# Scanning the rice Global MAGIC population for dynamic genetic control of seed traits under vegetative drought

**DOI:** 10.1101/2020.12.07.414474

**Authors:** Annarita Marrano, Brook T. Moyers

## Abstract

Grain size and weight are important yield components in rice (*Oryza sativa* L.). There is still uncertainty about the genetic control of these traits under drought stress, the most pressing emerging issue in many rice cultivation areas. To address this lack of knowledge, we investigated the genetic architecture of seed size, shape, and weight using the rice Global Multi-parent Advanced Generation Intercross (MAGIC) population, grown under well-watered and vegetative drought conditions. We measured variation in seed size and shape with a new high-throughput phenotyping method based on a desktop scanner and the open-source package Plant Computer Vision (PlantCV). Besides being affordable, rapid, and accurate, our method captured the phenotypic divergence between drought and well-watered samples, expressed as 12 different traits that include traditional size metrics and new grain shape measures. Overall, under water deficit, the MAGIC lines produced smaller and shorter seeds. We identified ten MAGIC lines with traits that make them good candidates for the release of rice cultivars with high yield potential under vegetative drought stress. We ran a marker-trait association analysis for the measured seed-related traits. Most of the identified marker-trait associations showed strong genotype-by-environment interactions (GxE), with most allele effects being conditionally neutral. These results suggest dynamic genetic control of seed size, shape, and weight under vegetative drought stress in rice, highlighting the importance of understanding the contribution of GxE interactions on trait variation to develop resilient and high-yielding rice varieties. Our study confirms that combining low-cost and high-throughput phenotyping strategies with a diverse genetic material suited for multi-environmental trial provides solutions for adapting rice cultivation to current and future environmental adversities.

## Introduction

Rice (*Oryza sativa* L.) is among the ten most cultivated crops worldwide, with 655.5 million tons produced in 2018 (FAOSTAT, 2018), and is the primary calorie source for more than half of the global population [1]. Large investments in rice genetic improvement began with the discovery and release of semi-dwarf, high-yielding varieties during the Green Revolution (GR) and yielded an almost three-fold increase in rice production in the decades following [2]. However, the current pace of yield increase is insufficient to meet future demands of a rising global population that is expected to reach almost 9 billion in the next 30 years [3].

While trying to boost yield rates further, rice breeders are dealing with the shortage of natural resources that is expected to become even more critical in the near future due to climate change [4]. Drought severity is the most pressing emerging issue in irrigated and rainfed rice production areas, where yield losses due to water shortage are already a reality [5]. The recent discovery of tight linkage between loci controlling plant height and yield under drought poses additional challenges for developing high yielding varieties under limited water conditions [6]. In particular, the strong selection and widespread adoption of semi-dwarf, high-yielding rice individuals during the GR era caused a loss of drought-tolerant alleles, leading to the high susceptibility to drought stress of most commercially grown modern varieties of Asian rice. The generation of cultivars carrying favorable alleles at loci controlling high-yield potential and drought resilience are promising strategies for a future, sustainable rice cultivation.

Seed (or grain) size and weight are two important components of rice yield. In addition, grain physical appearance, defined mostly by length, width, and shape, is a quality trait highly valued by consumers, whose preferences vary enormously across the world [7]. As most cereal seeds, a rice grain has its embryo and endosperm enclosed in a thin coat and covered by the spikelet hull (the husk), which defines the filling container. By setting the storage capacity of the grain, the husk has a crucial role in limiting grain growth and determining seed size [8]. Seeds grow through cell division and expansion, accompanied by nutrient accumulation and water intake. Drought stress seriously compromises grain filling by altering the source-sink balance necessary for carbon fixation and nutrient mobilization to the seeds. A reduced photosynthesis rate, leaf senescence, altered phloem turgor, and failure to form starch granules in the seeds are among the physiological responses to drought that cause short and incomplete seed filling [9].

Due to their importance as quality traits and yield components, grain size and weight have been amply studied in rice, and many major quantitative trait loci (QTL) have been identified for these traits [10–15]. However, the genetic control of seed-related traits under drought stress is still not fully understood. The laborious, time-consuming phenotyping procedures usually adopted to measure seed size and weight and the challenge of testing large, complex genetic populations under different environmental conditions are, most likely, the reasons for this slow progress.

High-throughput phenotyping of seed size and shape is incredibly labor-intensive, especially if executed using traditional methods based on manual measurements (*i.e.,* with calipers). In the last decades, the advent of digital phenotyping in agriculture has enabled automatic measurements of seed size and shape through the generation of several computer vision analysis tools [16], such as *SmartGrain* [17], *GrainScan* [18], and *PANorama* [19]. These methods identify single grains in seed images and acquire different size and shape metrics, including seed length, width, area, and perimeter. However, while these computational tools transformed the tiresome task of seed phenotyping through less labor-intensive and high-throughput procedures, they are either platform-dependent, not open-source, not easily parallelized, or not flexible to user preferences. Recently, Gehan *et al.* [20] released the latest version of the Plant Computer Vision (PlantCV) software package, an image processing toolkit designed in Python and comprising a collection of modular and flexible functions for plant image analysis. Besides being open-source and community-developed, PlantCV allows users to design versatile pipelines able to process different imaging systems (*e.g.,* visible, near-infrared, hyperspectral, etc.) and to investigate various plant features, including organ morphology and color variation [21].

Along with advances in phenotyping techniques for seed-related traits, rice genetics has seen incredible gene-trait discoveries in the last decade thanks to the development of strategically diverse genetic resources, such as Multi-parent Advanced Generation InterCross (MAGIC) populations [22]. A MAGIC population is obtained by intercrossing multiple parents, a design that allows sampling a greater proportion of genetic variation than the classical bi-parental populations while shuffling the genome and increasing recombination rates [23]. Therefore, MAGIC populations integrate the statistical power of bi-parental populations with the higher resolution of association panels. Among the many advantages of using multi-parent populations, two are particularly favorable for the study of drought effect on yield-related traits: (*i*) the segregation of multiple traits according to the selected parents; (*ii*) the generation of nearly completely homozygous lines by single seed descent, which enables experimental replication within environments and, therefore, the study of genotype-by-environment (GxE) interactions. In rice, MAGIC populations have been developed using the two major rice ecotypes, *indica* and *japonica* [22]. The founder lines are elite varieties or breeding lines with desirable profiles for traits essential across environment, such as yield, tolerance to abiotic stresses (*i.e.,* drought, salinity, and submergence), and disease resistance. One of the developed MAGIC populations is the “Global MAGIC”, obtained by crossing the 8-way crosses of elite *indica* and *japonica* MAGIC populations (for 16 parents in total). The Global MAGIC is, therefore, representative of the genetic background of both rice ecotypes, grown globally under different environmental conditions. *Japonica* rice is usually cultivated in temperate environments at high latitudes or altitudes, whereas *indica* rice is mainly grown in tropical and subtropical environments at lower latitudes or altitudes. Even though large variation in grain size and shape can be found within the single subspecies, typical *japonica* varieties produce round and medium/short grains, while *indica* cultivars tend to form long and thinner seeds [24]. This phenotypic divergence in seed size and shape between the two rice ecotypes derives from natural genetic variation at major QTL controlling grain length and width, such as *GS3* [25], *qSW5/GW5* [26], and *GLW7/OsSLP13* [27].

In the present study, we investigate the genetic control of seed-related traits under vegetative drought stress using the Global MAGIC population. We implement a new phenotyping procedure based on a desktop scanner and a user-defined PlantCV pipeline. This new method rapidly and accurately quantifies the morphological differences in seed size and shape, providing new measures of grain appearance, a trait that influences consumer preferences and defines market prices. We found environment-specific marker-trait associations, suggesting that (*i*) grain-related traits present different genetic architectures under drought and well-watered conditions, and (*ii*) GxE interactions play a crucial role in rice seed trait stability across environments.

## Methods

### Plant Material and drought treatment

Our analyses include 245 recombinant inbred lines from the Global MAGIC population developed at the International Rice Research Institute (IRRI) [22]. We planted seeds from the S6 generation using dry direct seeding at IRRI (14° 10’ N, 121° 15’ E) on 27-Jan-2015 (local dry season) in two environments: (*i*) under seedling stage drought conditions (DS), and (*ii*) under flooded, well-watered conditions (WW). We planted in an augmented design with the 16 parents, 10 MAGIC lines, and five standard IRRI checks replicated, and the remaining MAGIC lines unreplicated within four blocks.

We mechanically sowed seeds by placing one seed at a depth of 1–2 cm every 5.4 cm in plots of six 1.6 m rows with 20 cm row spacing in an open field. We planted the WW treatment on a lower terrace than the DS treatment. We sowed into dry, rotovated soil and established plant growth by sprinkler irrigation 2–3 times per week or moving the rainout shelter to allow rainfall to reach the plot.

We performed the DS treatment by stopping irrigation at 34 days after sowing (DAS). We monitored the level of drought stress using tensiometers installed at 15 cm and 30 cm depths. Throughout the DS treatment, we resumed irrigation at 45, 77, and 90 DAS based on the progression of the seedling stage drought. Only 117 mm of rain fell across the entire DS treatment. We flooded the WW environment at 34 DAS and maintained it flooded throughout the rest of the season.

At maturity, we hand-harvested and dried the panicles at 50 °C for three days. Then, we threshed and blew the dried panicles before determining grain moisture content, normalized to ca. 14%.

### Phenotyping

We measured seed size and shape using a desktop scanner (Epson Perfection V600 Photo Scanner, Epson America) and a user-defined pipeline designed with the PlantCV software package v3.0 [20]. In detail, we poured sample seeds on the desktop scanner in the middle of a phenotyping set-up consisting of a black tray, two color standards, and a ruler as a size standard (**Fig 1**). We built this simple setup to minimize variations of image size and light intensity across the scans. We saved images using prefixes, including phenotyping date (format MM-DD-YYYY), the scanner model, and a three-digit number indicating sample phenotyping order (*e.g.,* the first phenotyped sample was saved as 03-26-2020_EpsonV600_001.jpg). This prefix structure allows the PlantCV pipeline to read and process all images.

**Fig 1.**
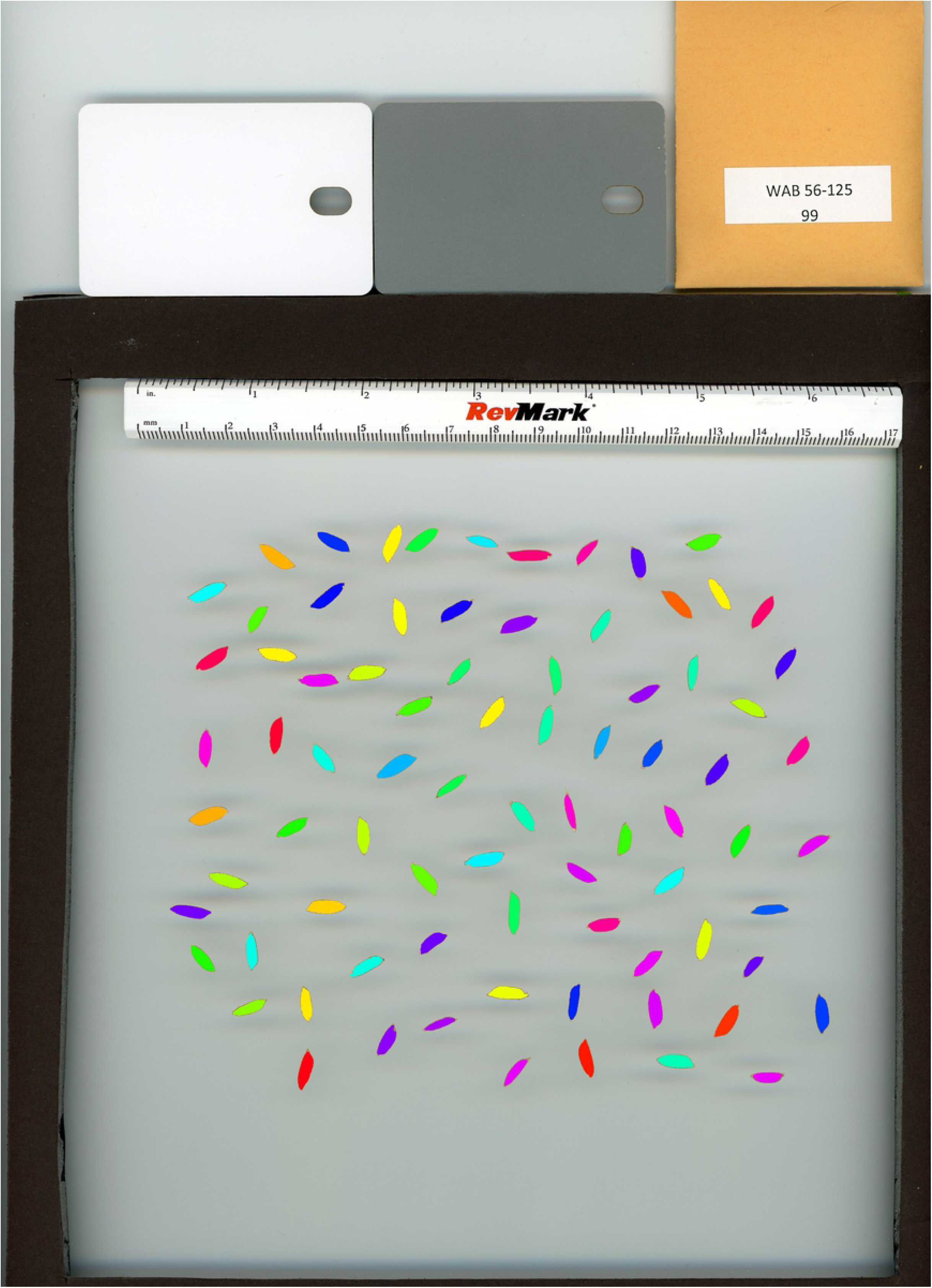
Seed image processing using a user-defined pipeline in PlantCV. After manual thresholding a seed image, the pipeline identifies and measures each seed separately. In the figure, it is also represented the simple set-up used for phenotyping rice seeds using a desktop scanner. The set-up includes a black tray to control image size variation, two color standards (white and grey cards) for image exposure normalization, and a ruler as a size standard.

We processed the RGB (Red Green Blue) images generated with the scanner using a personal laptop with Intel® Core™ i7 8650u CPU @1.90Ghz and 16 GB RAM. The PlantCV code used for this manuscript is available at https://doi.org/10.5281/zenodo.4156942, and more details on the PlantCV functions used in our pipeline can be found in the online user manual of PlantCV (https://plantcv.readthedocs.io/en/latest/). Briefly, for each RGB image, the pipeline first standardizes image exposure using the white standard color. Then, it separates the seeds from the background by applying threshold values in the LAB (L * a *b; where L = lightness, a = green/magenta, b = blue/yellow) color-space by so creating a binary image (mask) with white pixels as objects. Afterward, small holes in the objects are filled, and morphological filters are applied (erosion and dilation) to remove isolated noise pixels (*e.g.,* awns if present), define object boundaries and edges, and, therefore, separate seeds in close proximity or touching each other. This filtered binary image is then used to mask the normalized RGB image, and objects within a defined region of interest are detected and analyzed for shape characteristics (**Fig S1**). The 12 traits measured per seed are described in **Table S1**. We then parallelized the pipeline over all sample images as described at https://plantcv.readthedocs.io/en/stable/pipeline_parallel/. All trait estimates per seed and per sample are saved in JSON text files, which are then merged and converted to a final CSV table file using the accessory tool “plantcv-utils.py” implemented in PlantCV. All seed images are available at https://doi.org/10.5281/zenodo.4158169.

We also measured the grain weight of 50 seeds per sample using an analytical scale (Adventurer® Analytical, Ohaus, USA). We then converted the weight of grains to 1000-seed weight for easy comparisons with previous studies.

### Statistical analyses of phenotypic data

We performed all statistical analyses of the phenotypic data using the R statistical software version 3.5.2 [28]. For a matter of comparison with previous studies, we converted all traits on a pixel scale (**Table S1**) to millimeters (mm) by multiplying for 0.085, which is the conversion factor pixel-to-mm for an image with a 300-dpi resolution. We first removed outlier measurements by identifying seeds with area estimates either too large (*i.e.,* multiple adjacent seeds during the scan detected as a single object) or too small (multiple objects identified in one seed, which likely was fragmented in small pieces). We also obtained trait distribution and checked trait normality based on q-q plots generated with the function ‘qqnorm’ implemented in R.

We then estimated the significance of phenotypic differences among genotypes and treatments (WW vs. DS) by running the analysis of variance (ANOVA) test for the normally distributed traits and the Kruskal-Wallis test for non-Gaussian traits. We estimated individual median values per each seed size-related trait in each treatment to account for the differences in the number of seeds per sample, which were then used as phenotypic entry per genotype for all following analyses. We conducted a principal component analysis (PCA) for all traits in drought and well-water conditions using the ‘prcomp’ function implemented in R. We also computed trait correlations in each environment using the ‘rcorr’ function from the R package ‘Hmisc’ [29].

### Genotyping

We used the genotyping data generated for the Global MAGIC lines (S6 generation) using Genotyping-by-Sequencing (GBS) [30] as described in [31]. Briefly, we built GBS libraries with each Global MAGIC line and six biological replicates of each parent. We then sequenced the GBS libraries across 14 lanes with single-read Illumina sequencing technology. After, we processed the GBS data with the TASSEL 5.0 GBS v2 pipeline [32] to call variants against the *japonica* rice genome Nipponbarre (assembly IRGSP-1.0/MSU7) [33] and the *indica* genome 93-11 (assembly ASM465v1 from BGI) [34]. In detail, we first created 917,254 tags using the GBSSeqToTagDBPlugin (minimum kmer count 100, kmer length 75, minimum kmer length 20, minimum quality score 20, maximum kmers in database 100M) and then mapped them onto each reference genome with Bowtie2-2.2.6. We identified single nucleotide polymorphisms (SNPs) and indels in the aligned tags with the DiscoverySNPCallerPlugin, and scored them for coverage, depth, and genotypic statistics using the SNPQualityProfilerPlugin. Using statistics from the genotypes of the six biological replicates of the 16 parent lines, we removed variants with less than 50% coverage, heterozygous calls, more than two alleles, and inconsistent genotype calls for all replicates of a parental line. We finally retained sites with genotype calls for at least 15 out of the 16 parents with the biological replicates merged. We filtered out Global MAGIC lines with more than 90% missing data.

### Genome-wide association and candidate genes for seed size and shape

We carried out a marker-trait association analysis using a general linear model (GLM) approach implemented in TASSEL 5. Since we did not observe significant population structure within our Global MAGIC population (**Fig S2**; [31]), the GLMs did not include any covariates to account for underlying structure of population. We ran GLMs for all traits in each environment and corrected for multiple testing using an FDR (False Discovery Rate) ≤ 0.05 as significance threshold. We searched for known QTLs and candidate genes for drought and seed size-related traits around the identified marker-trait associations using rice public databases, including SNP-Seek II [35], RAP DB (Rice Annotation Project Database) [36], and Oryzabase [37]. We also investigated the genomic regions surrounding the most significant associations for additional candidate genes using the most recent gene annotation of the reference genome Nipponbare (MSUv7).

For assessing the genotypes at the 16 sites within the gene LOC_Os12g27810, we downloaded the whole-genome variant calls of the 3,010 Asian rice accessions against the Nipponbare reference as indexed VCF files from Amazon Public Data at https://registry.opendata.aws/3kricegenome/, [38]). We then filtered for the GBS variants discovered against Nipponbare in the Global MAGIC population using the commands *--position --recode* of VCFtools v0.1.15 [39]. We merged all filtered VCF files with the Perl tool vcf-merge implemented in VCFtools, after compressing (by *bgzip)* and indexing (by *tabix).*

## Results

### A new phenotyping pipeline for seed size and shape in rice

We phenotyped 535 seed samples in total, of which 263 were drought-treated and 272 from the well-watered environment. The phenotyping dataset included 14 of the parents (two are under restricted material transfer agreement at IRRI), of which Sambha Mahusuri + Sub1 and IR-77298-14-1-2-10 did not produce any seeds under drought conditions. On average, we scanned 110 seeds per sample in ca. five minutes, with a sample seed number ranging between three and 245. All images were then processed using a user-designed pipeline in PlantCV, which uses a manual thresholding method to segment the seeds in each image (**Fig 1; Fig S1**) and measures 12 different traits related to seed size and shape (**Table S1**). The PlantCV pipeline took about 6 hours to upload and analyze all seed images simultaneously on a standard laptop. Therefore, assuming flexible working time for phenotyping and the efficiency of the PlantCV pipeline to process hundreds of images at the same time, our method allows screening seed size in large populations in less than a month.

Our method captured the phenotypic divergence between samples grown under well-watered and drought conditions (**Fig 2**), providing new measures of seed size and shape in addition to traits as seed length and width widely implemented in rice. For instance, ellipse eccentricity (**Table S1**) is a measure of the roundness of an object, with values ranging from 0 (perfect circle) to 1 (elliptic shape). Another seed shape measure is solidity, which is the ratio between seed area and convex hull area, where a convex hull is defined as the smallest convex boundary containing a shape or a group of points. Values of solidity lower than one indicate seeds with an unusual or aberrant shape that need a bigger convex hull to contain them.

**Fig 2.**
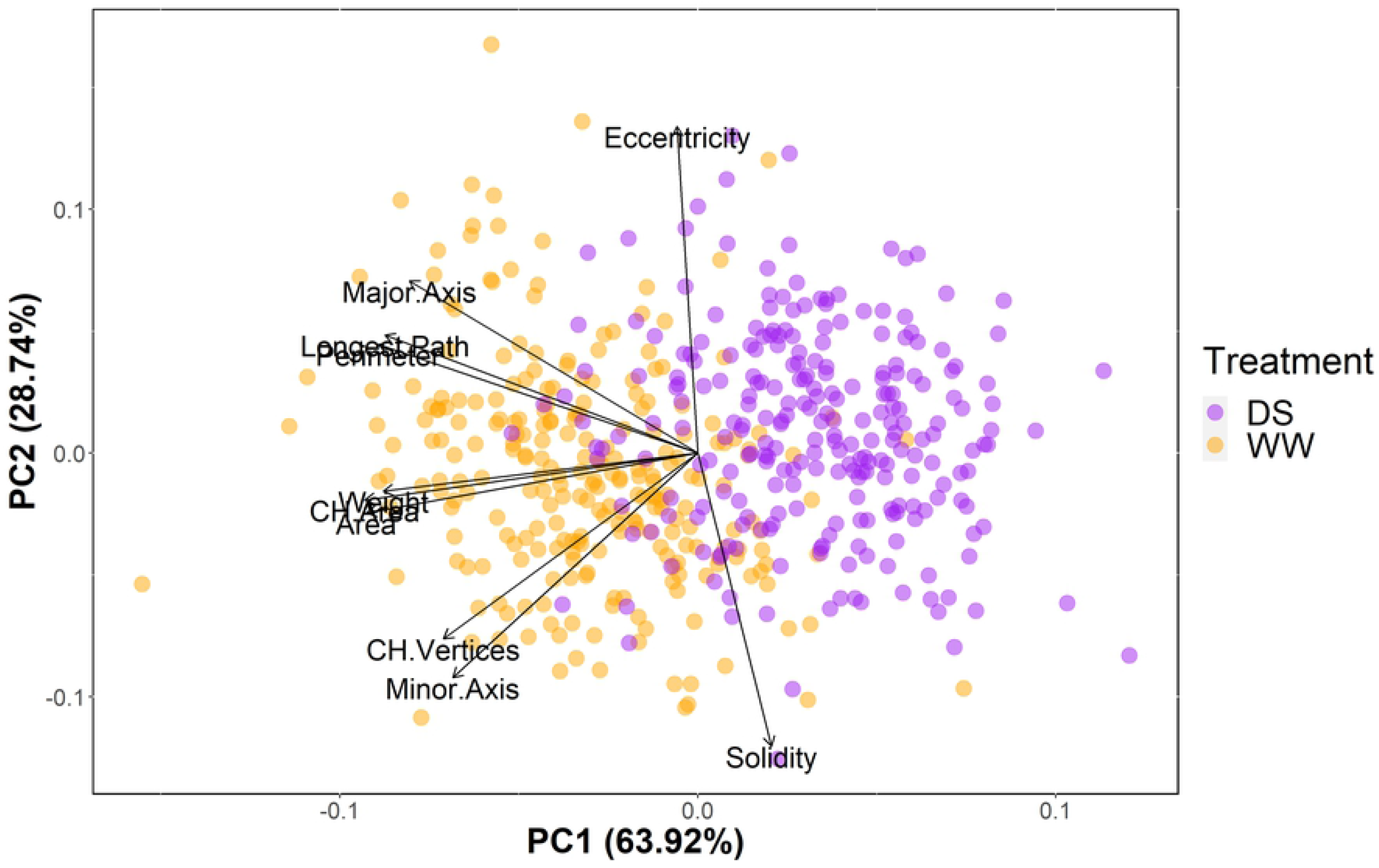
Projection of individual trait phenotypes on the first two axes of the Principal Component Analysis (PCA). Loadings are also reported for ellipse eccentricity, ellipse major and minor axes, longest path, perimeter, seed weight, convex hull area (CH.Area), convex hull vertices (CH.Vertices), seed area, and solidity. DS = drought stress; WW = well-watered.

To evaluate the accuracy and reproducibility of our phenotyping method for seed size in rice, we re-measured 107 samples selected randomly among the original 535. The re-phenotyping dataset included 53 DS and 54 WW seed samples, with a mean number of 112 seeds per scan. We then compared these new trait measurements with those observed during the first phenotyping for the same samples (**Fig 3**). Overall, the coefficient of correlation between the first and second phenotyping was higher than 0.9 for all traits but ellipse angle (R = 0.1; p-value = 0.31) and width (R = 0.74; p-value < 2.2e^-16^). In particular, width accuracy further decreased when considering DS (R = 0.64; p-value < 2.2e^-07^; **Fig S3**) and WW (R = 0.48; p-value < 0.00025; **Fig S4**) samples separately. The PlantCV function *‘analyze_object’* used in our pipeline defines width and length as the total span of object pixels along the *x*-axis and the *y*-axis, respectively [21]. Therefore, depending on seed orientation on the scanner, width, and length can change considerably across different scans of the same individual (**Fig S5**).

**Fig 3.**
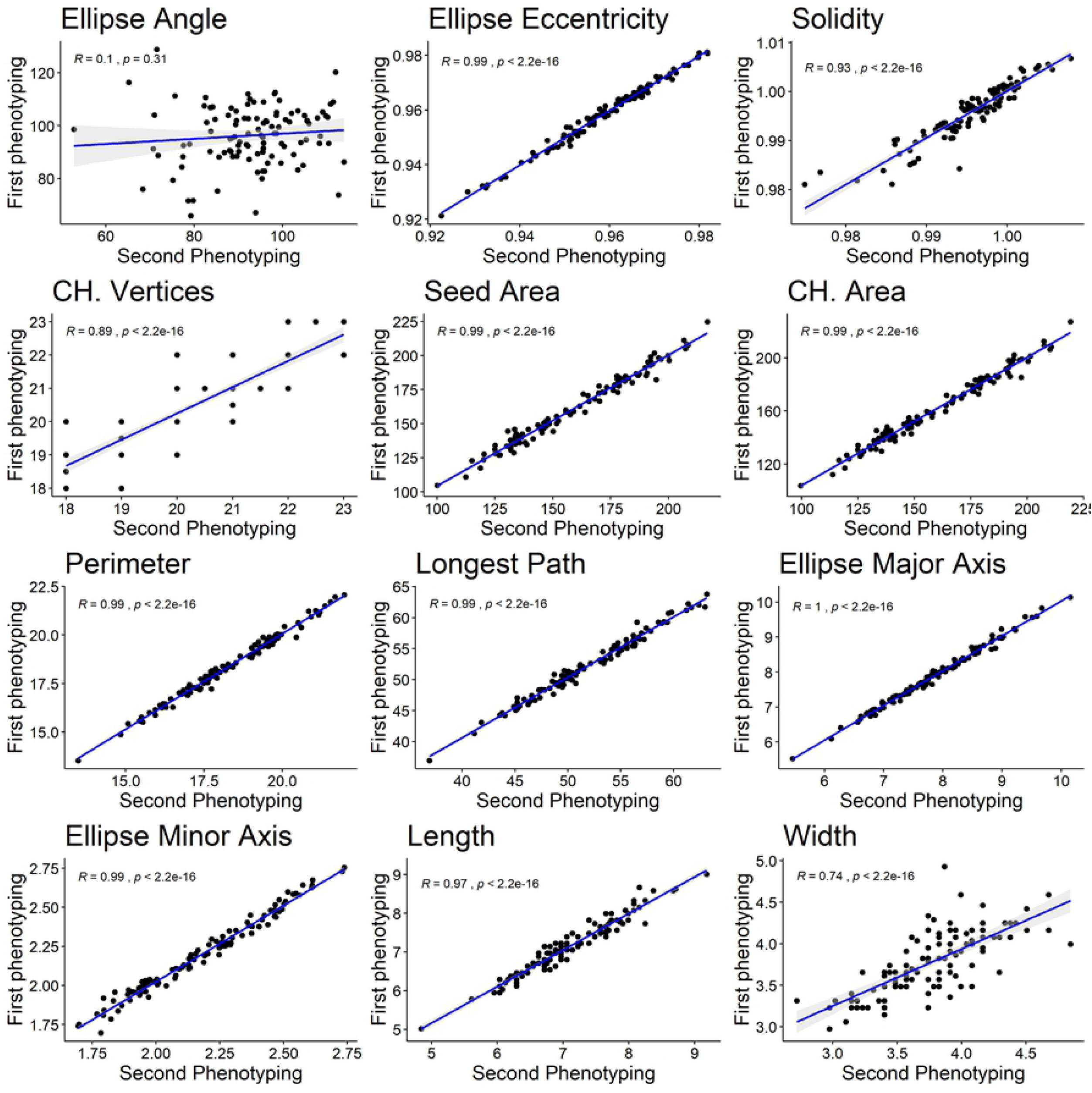
Correlation between trait measures of the first and second phenotyping of the 107 samples chosen randomly within both the drought and well-watered experiments. CH = convex hull.

On the contrary, ellipse major and minor axes are, by definition (**Table S1**), more accurate estimates of seed length and width, respectively. Seed orientation also explains the low consistency of the ellipse angle across measurements; in fact, the ellipse angle is the rotation angle of the bounding ellipse major axis (**Fig S5**). Our findings suggested that, while width, length, and ellipse angle can be useful measurements of seed size and shape for other crops, they are not suitable for rice grains unless seeds are deliberately constrained to a fixed orientation. Therefore, we decided to exclude them from our following analyses. On the other hand, we can conclude that our method is highly accurate and reproducible for the remaining traits.

### Seed-related traits under drought in rice

All measured traits exhibited a normal distribution, except for solidity and eccentricity (**Fig S6-7**). For all traits, we observed significant differences among individuals and treatments (p-values < 0–0.001; **Tables S2-S3**), suggesting both genetic and environmental control of seed size and shape in rice. Seeds from the DS and WW treatments are strongly differentiated along the first phenotypic principal component (PC1), accounting for more than 50% of sample phenotypic variance (**Fig 2**). The traits loading most heavily on PC1 are seed area, convex hull area, ellipse major axis, perimeter, longest path, and seed weight (**Fig 2; Table S4**).

We observe strong correlations between common and newly defined seed size traits and shape (**Fig 4**), *e.g.,* seed perimeter and longest path, or seed area and convex hull area. We see similar trait correlations in well-watered and drought-treated samples (**Fig 4**). Interestingly, ellipse minor axis is strongly negatively correlated with ellipse eccentricity, suggesting that seed shape depends on seed breadth. Ellipse major axis, seed perimeter, and longest path are all positively correlated with ellipse eccentricity. In addition, ellipse major and minor axes and seed area are positively related so that larger seeds are usually also longer or wider. As expected, grain weight correlates positively with almost all seed size traits, particularly with seed and convex hull areas.

**Fig 4.**
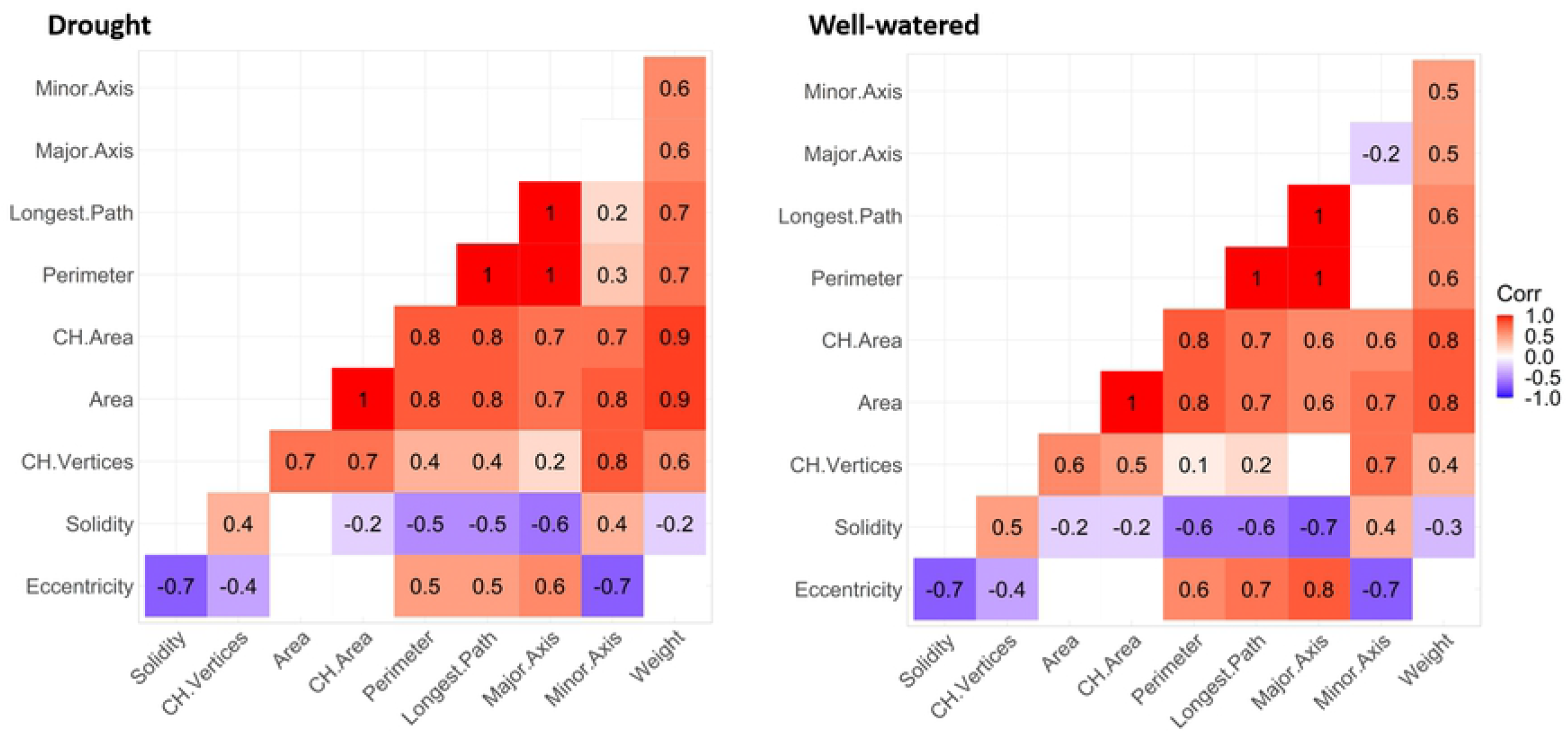
Pearson’s correlation analysis among the ten measured traits for the drought and well-watered samples. Blank squares indicate no significant correlations (p-values > 0.05).

Generally, seeds under drought were smaller and shorter than the well-watered, which, by contrast, showed more variability among genotypes (higher standard deviations; **Table S4**). However, 11 samples either outperformed under drought stress, producing seeds with bigger area, or yielded seeds of similar size besides the water availability condition during the growth (**Fig 5**; **Table S5**). The high yielding and blast resistant *indica* variety Shan-Huang Zhan-2 [40] was the only MAGIC parent among the top-ranked samples for seed area (**Fig 5**); its seeds had longer ellipse major axis and perimeter and a slightly bigger area under drought stress than well-water conditions (**Fig S8-S9**). All the remaining top-ranked samples were MAGIC lines, with MG-7741 being the best performing sample under drought stress; it produced bigger seeds with longer ellipse minor axis (**Table S5**). Eight samples of the top-ranked for seed area under drought also shown higher or similar values of grain weight, including the *indica* parent Shan-Huang Zhan-2.

**Fig 5.**
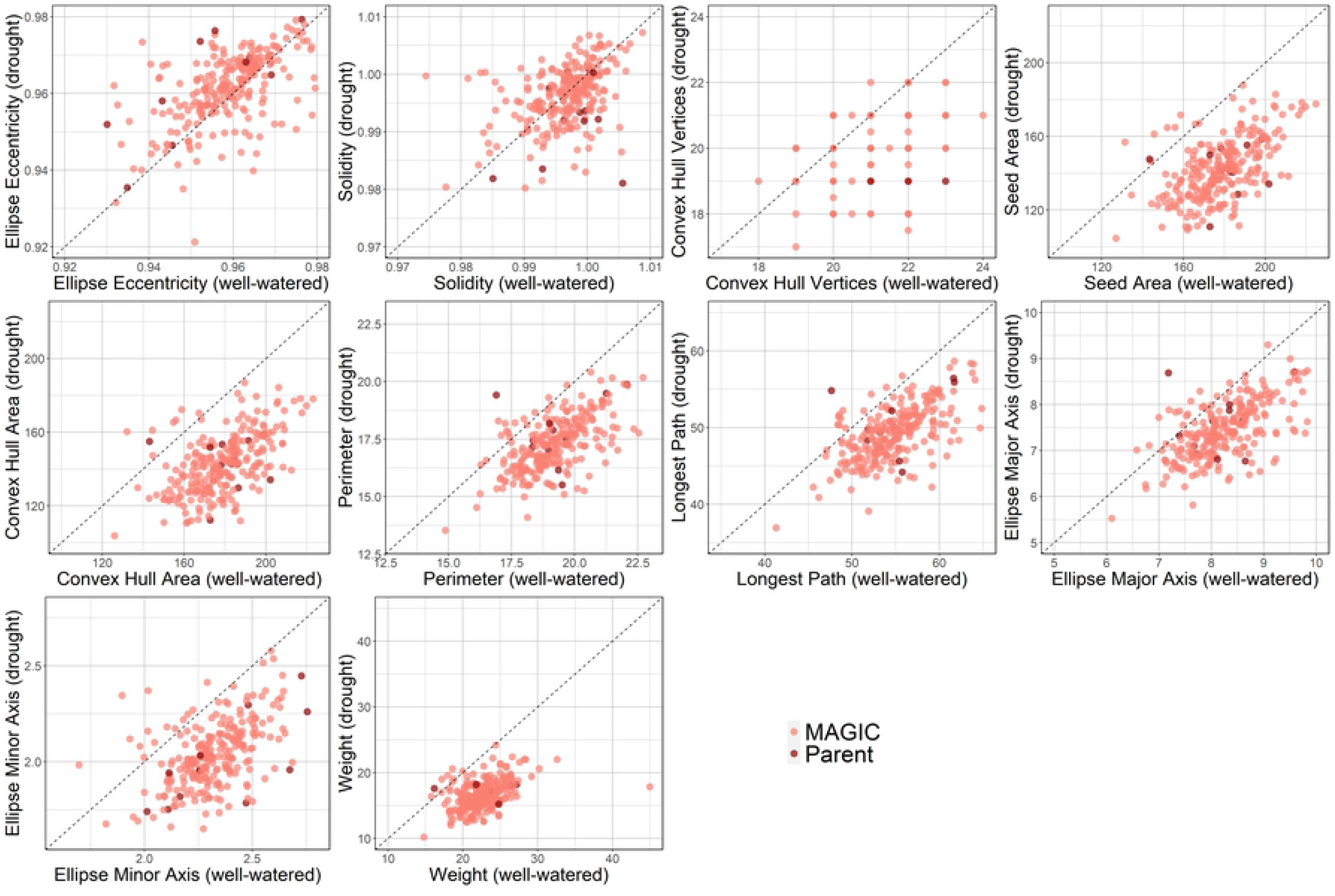
Individual trait values under drought and well-watered conditions. The dots in the left-upper triangle indicate individuals with higher trait values under water deficit. For instance, considering seed area, 11 individuals produced bigger seeds or grains of the same size under drought conditions; the parent Shan Huang Zhan-2 is among these individuals (the only dark red dot above the diagonal in this panel).

### Genetic architecture of seed traits under drought

In total, we identified 276 significant marker-trait associations across both environments and both genomic variant datasets (138 on Nipponbare; 138 on 93-11; **Fig S10-S11**). The two datasets are largely overlapping, but we see associations in each genome with sites that are not present in the other genome. We summarize the GWAS results in **Table S6**, where we organized them in intervals based on their physical vicinity and overlap across genomes. The full and detailed lists of significant sites are in **Table S7-S8**.

Under the WW environment, seed weight exhibited multiple trait-loci associations, confirming its quantitative nature. In particular, we detected genomic regions involved in weight control under normal irrigation conditions on the chromosomes 1, 3, 4, 5, 6, 7, 8, and 12 in both reference genomes and on chromosome 9 of 93-11 only (**Fig 6-S11**). In addition to seed weight, we found significant associations on chromosome 6 and 10 (Nipponbare only; **Fig 6**) in the WW environment for the trait solidity, a measure of seed shape.

**Fig 6.**
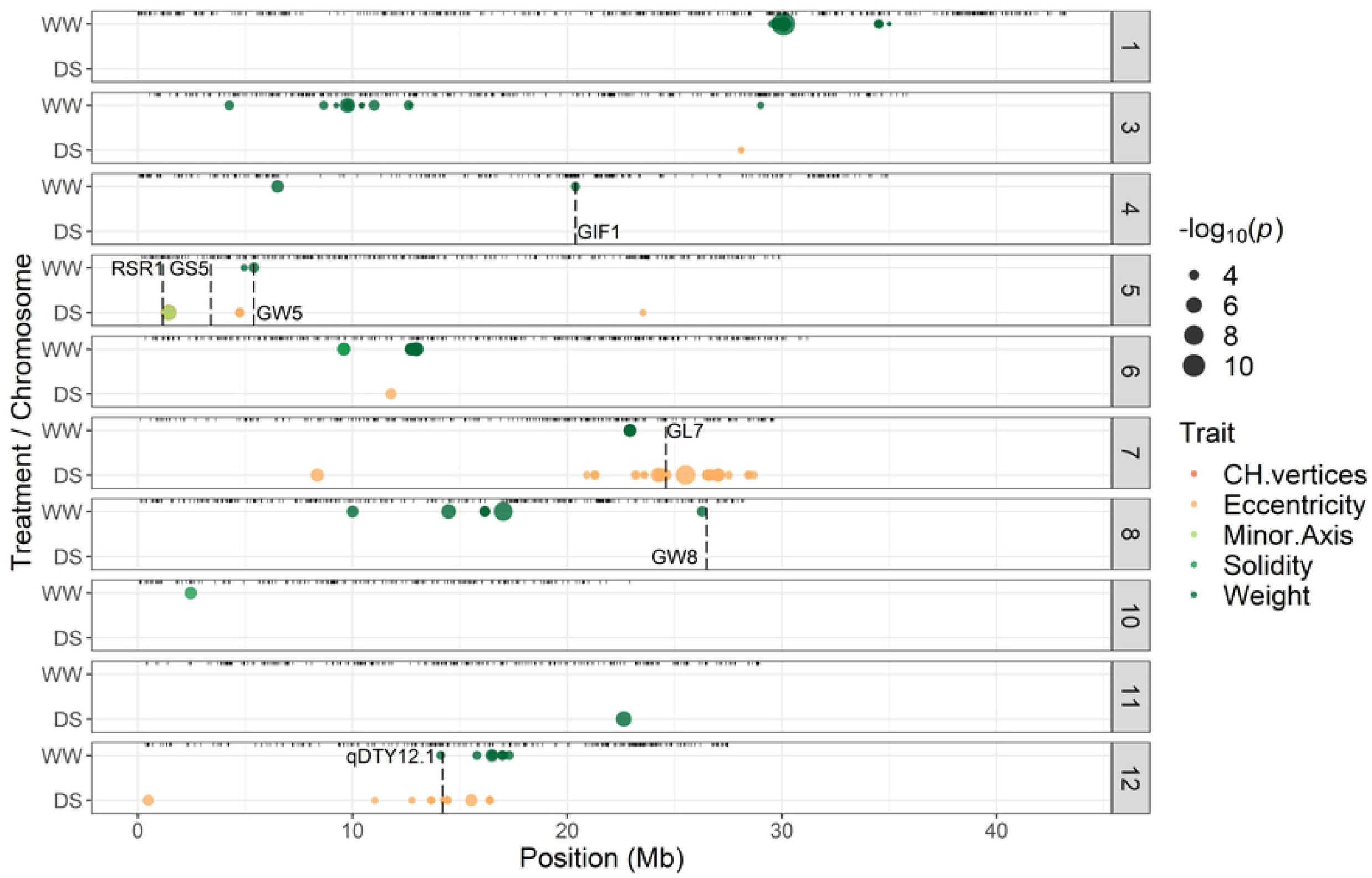
Genome-wide associations (FDR ≤ 0.05) found on the Nipponbare genome. Only chromosomes and traits with significant associations in at least one environment are shown. The x-axis reports site location (Mb) per each chromosome (vertical panel), while the y-axis indicates the environment specificity of the significant association (WW= well-watered; DS = drought stress). Circles are colored according to the associated trait, and their size is proportional to -log10(uncorrected p-value).

We detected significant associations in the DS environment, mostly for eccentricity on chromosomes 3, 5, 6, 7, and 12 of both subspecies’ genomes. We detected additional peaks for eccentricity on chromosomes 2 and 9 of the 93-11 genome and on chromosome 11 in the Nipponbare genome (**Fig 6-S11**). One site (S5_1439326 and S5_1530423 in the Nipponbare and 93-11 variant datasets, respectively) is significantly associated with eccentricity, convex hull vertices, and ellipse minor axis under drought stress. Based on the correlation indices, higher values of ellipse minor axis, which is a measure of seed breath, are typical of elliptic and less rounded seeds, whose convex hull would have more vertices.

We observe almost all associations in only one environment, indicative of different allelic effects depending on the environmental context (GxE). To explore this pattern, we evaluated the GxE allelic effects using trait mean in each environment as an approximation of environmental differences affecting that trait (**Fig 7-S12**). We notice changes in allelic effect sizes for seed weight, solidity, and eccentricity under drought and well-watered conditions (**Fig 7**). Most all alleles have large effects in one environment that decrease towards smaller or neutrality (no effect) in the other. A few alleles, though, present an antagonistic effect on seed weight across the two environments, causing a positive increase of grain weight under drought and a reduction of ca. 3 STD in the WW environment (**Table S7**).

**Fig 7.**
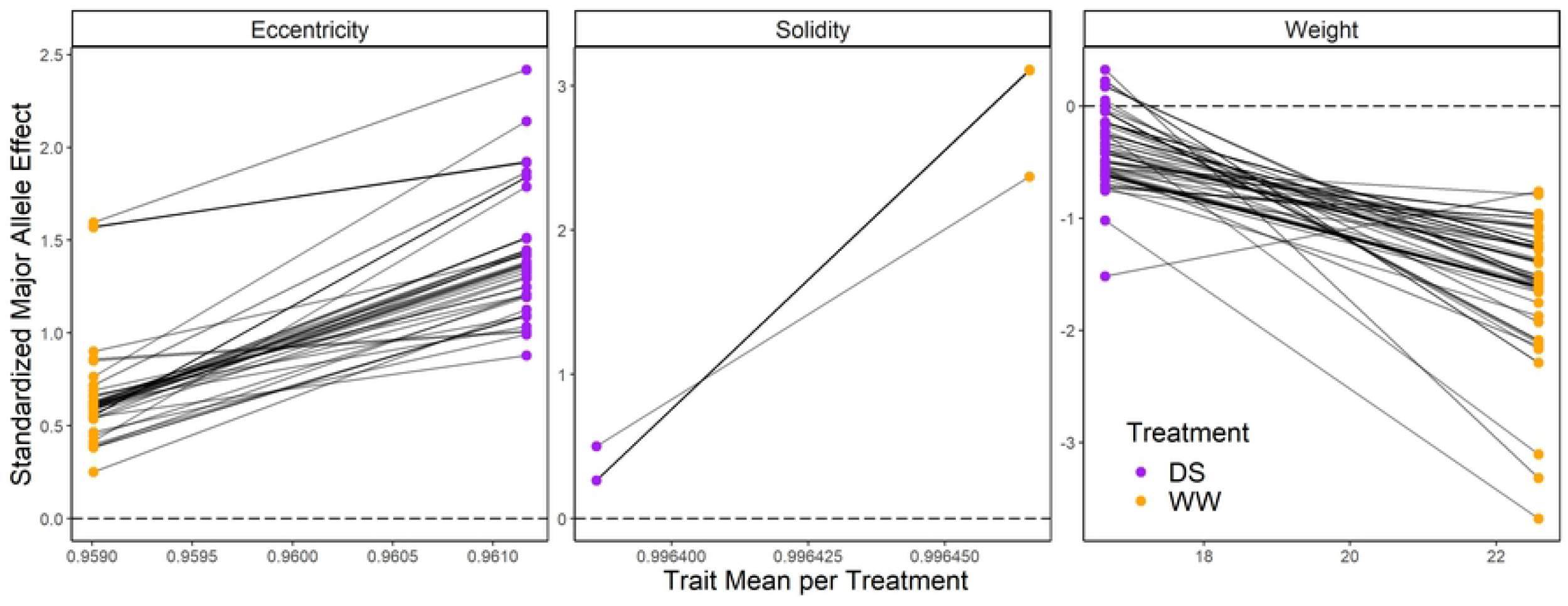
Reaction norms of the standardized allele effects for all Nipponbare sites significantly associated (FDR ≤ 0.05) to ellipse eccentricity, solidity, and seed weight in at least one environment (or treatment). Major allele trait effects standardized to trait standard deviation in each environment (y-axis) are plotted against the trait mean in each environment for the 243 lines common to the two environments (x-axis).

GxE interactions are less intuitive for the single site at 1.44 Mb on chromosome 5 associated simultaneously with convex hull vertices, ellipse eccentricity, and ellipse minor axis. Here we observe a consistent allelic effect for minor axis and convex hull vertices across the two environments that inverts direction for eccentricity, where the major allele has a larger effect under drought (**Fig S13**). Therefore, the major allele causes slender and more elliptic seeds under water deficit. These patterns of allelic effect size reflect the direction of the correlation among traits (**Fig 4**) that is, most likely, pleiotropic.

### Known QTLs and candidate genes around the significant associations

We examined the presence of QTL or candidate genes already identified for seed size and drought responses in rice, within intervals of 4 Mb around the association signals (2 Mb downstream and upstream the variant physical position). We defined window size according to the linkage disequilibrium (LD) decay of ca. 2Mb recently reported for the rice Global MAGIC population by [31]. All the results here reported refer to the marker-trait associations identified within the Nipponbare variant dataset due to the broader use of the *japonica* reference genome in previous genetic studies and the availability of an accurate and updated gene annotation.

We identified 68 QTL (**Table S9**), and 54 trait candidate genes (**Table S10-S11**) previously identified for seed and drought-related traits. For instance, the single marker-trait association for seed weight on chromosome 4 is 34.4 kb downstream of the GRAIN INCOMPLETE FILLING 1 (*GIF1)* gene, a potential domestication gene involved in carbon partitioning during early grain-filling [41]. Also, three know gene loci involved in seed size and shape (*i.e.,* RICE STARCH REGULATOR 1-*RSR1;* GRAIN SIZE 5 – *GS5;* GRAIN WIDTH 5 – *GW5*) mapped closely to the significant associations found on chromosome 5 (**Fig 6**). Other important gene loci co-mapping with the found trait associations are the GRAIN LENGTH (*GL7)* gene on chromosome 7 and the GRAIN WIDTH (*GW8)* gene on chromosome 8.

We additionally search for new candidate genes by exploring the Nipponbare genic space around the identified significant associations. We found 15,374 genes in the designed 4 Mb windows (**Table S12**). One of our genotyping-by-sequencing tags happens to span both a SNP and a 15-bp indel in the gene LOC_Os12g27810. This locus, which encodes an F-box domain containing protein, is upregulated under drought and it is located under *qDTY_12.1_* (15.8-17.4 Mb), a major QTL controlling grain yield under reproductive drought stress (**Fig 6; Table S9**) [42]. The SNP by itself would be nonsynonymous (a glycine to cysteine substitution at amino acid residue 357), but in combination with the indel creates the greater physicochemical change of a tryptophan substitution at that residue followed by a loss of five amino acids (residues 358 to 362). The indel and the SNP are separated by a single base pair and so are in strong linkage disequilibrium (*r^2^* = 1 in our data), forming two haplotypes (haplotype 1 = GGCGGCGCCAGCGCCAG; haplotype 2 = TG---------------). Haplotype 2, inherited from parent Jinbubyeo and present in only four of our 245 phenotyped MAGIC lines, is strongly associated with rounder seeds, particularly in the drought environment (**Fig 8**).We investigated the variation at the gene LOC_Os12g27810 in the 3,010 (3K) Asian rice accessions recently whole-genome sequenced by [38]. The 16 sites found associated with ellipse eccentricity in our analysis formed 24 distinct haplotypes in the 3K rice panel, of which none include the deletion, and only seven are present in more than one accession (**Table S13**). The haplotype 3k.Hap1 (the same as haplotype 1 in the MAGIC population) is the most abundant in the entire collection in general, even if over 70% of the accessions carrying it are *indica* (**Table S14**). The second most frequent haplotype is 3k.Hap2, which has our minor allele at the SNP locus S12_16392557, so the glycine to cysteine substitution, but not the deletion we saw in our population. This haplotype is present mostly in temperate (GJ-tmp; 54%) and subtropical (GJ-sbtrp; 21%) *japonica* accessions (**Fig 9**). Finally, the third most abundant haplotype has missing calls to all sites, which might be due to low coverage or low-quality sequencing data at this locus in the 221 accessions carrying the 3k.Hap3, or bioinformatic difficulties with coverage at these sites due to a segregating deletion. We suspect the latter scenario is likely, as over 50% of the individuals with missing calls (or 3k.Hap3) are tropical *japonica* compared to only 12% of the 3k in total (**Table S14**). We would not expect low coverage or sequence quality to be biased by taxonomy in this dataset.

**Fig 8.**
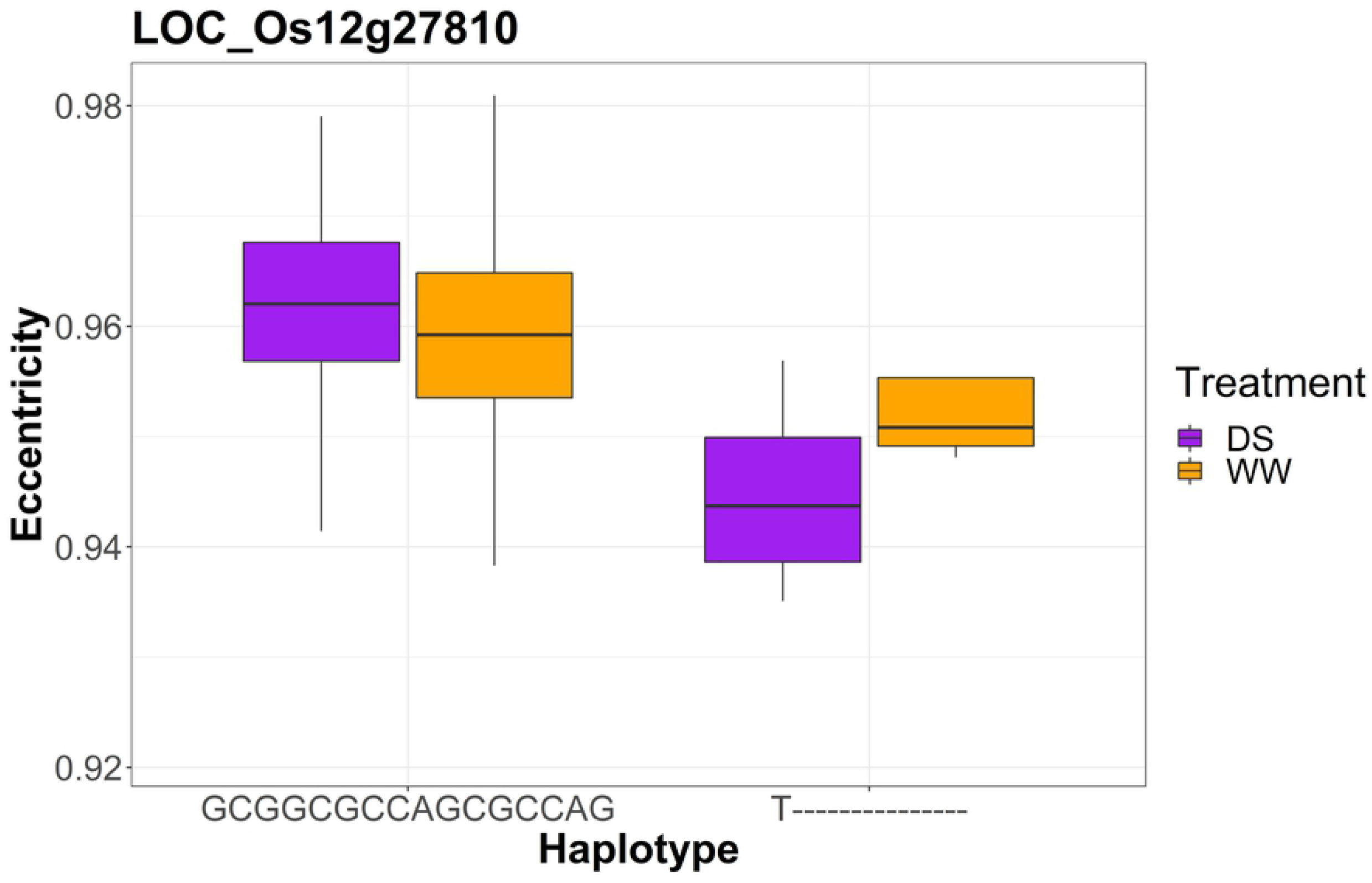
Phenotypic variance in the two environments (DS = drought stress; WW = well-watered) among the MAGIC lines carrying the two different haplotypes at the gene LOC_Os12g27810.

**Fig 9.**
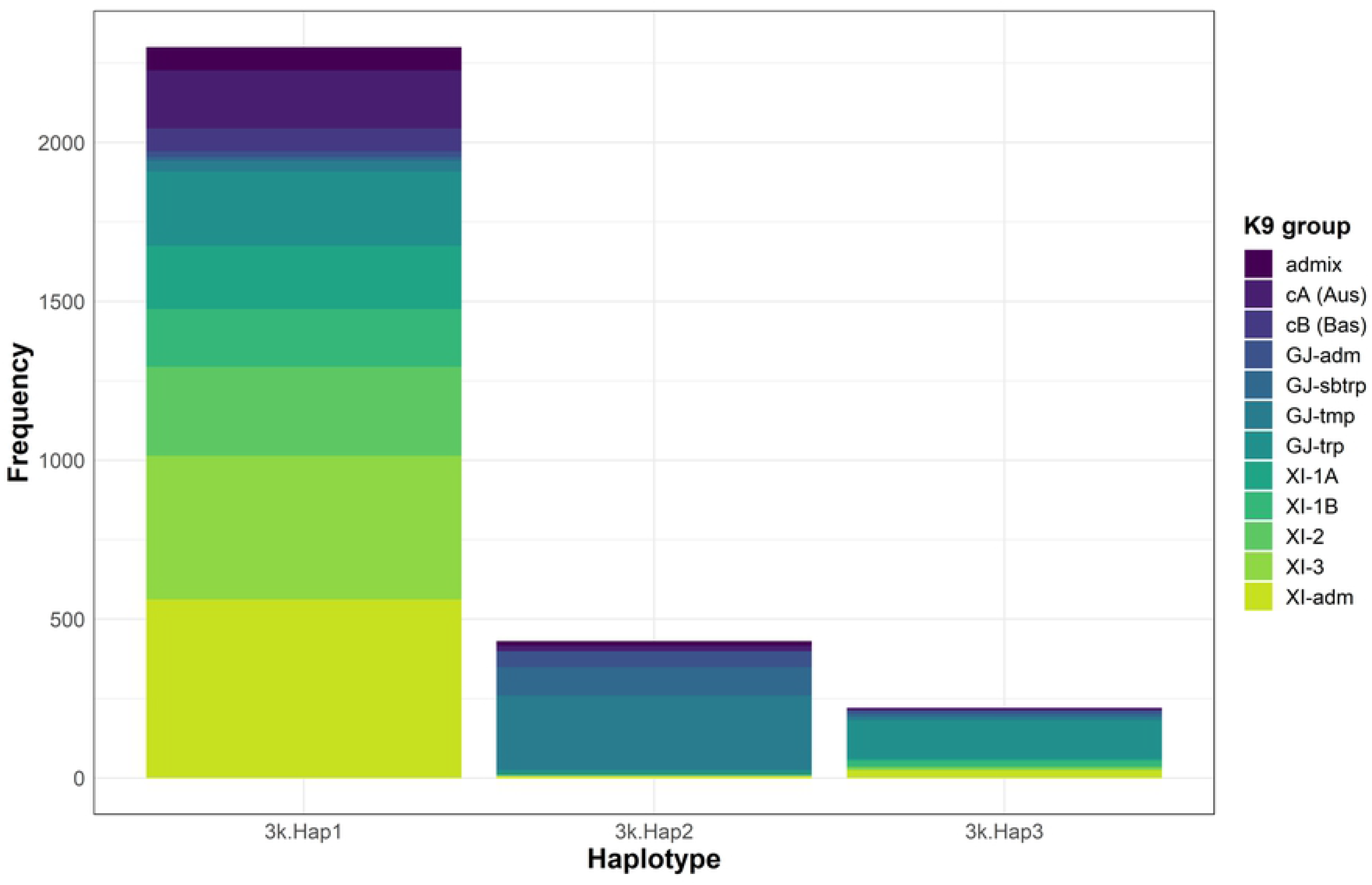
Frequency of the three most abundant haplotypes found at the locus LOC_Os12g27810 among the nine rice subpopulations identified by Wang et al. [38] after resequencing 3,010 Asian rice accessions. XI-1A = *indica* samples from East Asia; XI-1B = modern *indica* varieties of diverse origins; XI-2 = *indica* samples from South Asia; XI-3 = *indica* samples from Southeast Asia); GJ-tmp = East Asian temperate *japonica* samples; GJ-sbtrp = *japonica* accessions; GJ-trp = Southeast Asian tropical *japonica* samples; cA = Aus varieties; cB = Basmati and Sadri aromatic varieties.

## Discussion

Beyond their agronomical importance for crop production, seed traits are related to plant ability to adapt and survive in unfavorable environments [43]. Generally, plants producing larger seeds have a higher tolerance to biotic and abiotic stresses, including drought [44], although there are potential constraints and trade-offs for seed size. We studied the impact of water shortage during seedling stage on seed size, shape, and weight in rice, whose production in many cultivation areas is already facing severe losses due to drought stress. Our study not only provides evidence of a different genetic architecture of seed-related traits under variable environmental conditions but also identifies possible donors for stable yield component traits under drought in rice. These individuals are resilient to limited water conditions during seedling vegetative growth, an adaptive capacity found to be correlated with large grain size [44]. These results are promising for future adoption of dry-direct seeding as a cropping system also in lowland rice, whose modern cultivation in fully irrigated fields demands high resource consumption, notably water [45].

To screen seed size and shape in over two hundred individuals grown in two environments, we needed a phenotypic method that was rapid, affordable, less labor-intensive, and, especially, accurate. Digital imaging is the answer to all these needs, with different software already available for automatic and high-throughput phenotyping of seed size and shape. In rice, *SmartGrain* [17] has been adopted to analyze seed images and measure grain size and shape in MAGIC populations [46,47]. However, *SmartGrain* works only on Windows platforms, and it does not allow to parallelize a pipeline across multiple images. Instead, we processed all seed images using PlantCV, which overcomes these limitations and in addition, allows one to easily integrate different types of image analysis (*e.g.,* morphology, color variation) into a single workflow. For instance, Tovar et al. [48] used PlantCV to measure seed and panicle size under heat stress in quinoa (*Chenopodium quinoa* Willd.). Our study confirms the eligibility of PlantCV as a reliable and flexible solution for seed size phenotyping.

Besides being less time-consuming and affordable, our method captured the grain morphological differences between rice lines grown under well-watered and drought conditions (**Fig 2**), providing new measures of seed size and shape, such as ellipse eccentricity and solidity, in addition to the standard grain length and width used in rice. By rescanning a subset of about 100 samples, we observed low consistency across measurements for ellipse angle and seed width, due to the variation of seed orientation on the scanner across experiments, and algorithm limitations in resolving these discrepancies. Repeating seed phenotyping was crucial to identify these inconsistencies that could have otherwise biased the following genetic analyses and the conclusions drawn upon them. In the midst of plant phenomics growth as main field in agriculture, and the consequent development of numerous platforms for image data analyses, our results highlight the importance of verifying the accuracy of the selected pipeline in quantifying and interpreting the biological diversity of the studied crop system. It can be all too easy to forge ahead with applying a new method without first verifying its accuracy: we caution against this.

We ran GWAS for nine traits related to seed size and shape, along with 1000-seed weight, identifying significant associations mostly for ellipse eccentricity in the drought environment and seed weight in the well-watered environment (WW). These results are potentially biased by the fact that only about half of the entire Global MAGIC population produced seeds under seedling stage drought stress, inevitably limiting our power to identify additional marker-trait associations for the analyzed seed traits. We could phenotype only 245 of the 523 MAGIC lines that were planted under both drought-stressed and well-watered conditions. On the other hand, genetic control of seed size and shape is irrelevant if seeds are never produced, and it is important to study genetic architectures in the appropriate context. We conducted our GWAS in the set of lines that produced seed under both conditions so that we could directly compare allelic effects across environments.

The two genomic variant datasets shared almost all significant associations, with a few exceptions that could indicate private structural variation (SV) within the *japonica* or *indica* subpopulations. As the Global MAGIC population has parents from both subpopulations as well as one from the smaller Basmati subpopulation, we could expect subpopulation specific SV to be segregating. The rice pan-genome analysis performed by Wang et al. [38] shown that the *indica* and *japonica* subspecies are different for about 14.9% of the genes and 14.3% of the gene families. For instance, we found a Nipponbare exclusive significant peak for seed weight under drought stress at 22.64 Mb of chromosome 11. This marker-trait association is about 7 Mb apart from an association signal for raw grain shape found only in *japonica* and included in a syntenic break on chromosome 11 between the two reference genomes Nipponbare and 93-11 [49]. These findings highlight the importance of genomic structural variation in the genetic control of trait variability between the two rice ecotypes, a potential source of divergence that needs to be further addressed in future genetic studies in rice.

Both ellipse eccentricity and grain weight depend on a regular and complete grain filling process, which starts with the onset of plant senescence and the remobilization of carbon reserves from source tissues (mainly leaf and stem) to grains [50]. In particular, senescence initiation activates starch degradation in the source tissues, followed by the mobilization of sucrose molecules to the grains, where it is used to form starch granules. Therefore, the amount of carbon fixed and accumulated in vegetative tissues, the efficiency of sucrose metabolism, and the starch-synthesizing capability of endosperm are among the main factors determining grain filling rate and duration.

In rice, water deficit during grain filling stage accelerates senescence and the remobilization of carbon reserves to the seeds, causing an increase grain filling rate and a reduction of grain filling duration [51]. Here, we applied drought stress during early vegetative growth, which could have affected sink strength and the formation of temporary starch reserves during the first phases of plant growth. Differences in photosynthetic rates and carbon fixation in response to drought could explain the variability in grain filling capacity and, thus, grain weight and shape within our MAGIC population. Consistently, we found marker-trait associations in the proximity of known loci involved in grain filling and seed starch metabolism. On chromosome 4, a significant association for seed weight is located 34.4 kb apart from the *GIF1* locus that encodes a cell-wall invertase required for carbon portioning during the early stages of grain filling [41].

In particular, *GIF1* has a crucial role in sugar unloading during grain development, and a 1-nt deletion in the coding region causes low sucrose levels in the seed, slow grain filling, chalkiness, and reduced grain weight. Another locus involved in the first stages of seed development is *RSR1,* found near (283.3 kb) the multi-trait association on chromosome 5 for eccentricity, solidity, and ellipse minor axis. *RSR1* negatively controls the expression of starch synthesis genes, and its deficiency causes enlarged seeds and increased seed weight, without any changes in seed starch content [52]. Therefore, the identification of *RSR1* as a candidate gene for seed size and shape in rice confirms the key role of sink strength in grain filling [53] and suggests that water shortage during vegetative growth might influence individual sink capacity with cascading effects on grain size and quality.

Among the known candidate loci controlling grain size and weight in rice, our GWAS peaks co-localize with *GW5, GL7, and GW8.* These major QTL have key roles in either cell division or elongation during grain development [24,54–56] and have been identified in previous genetic analysis of seed size and weight in *indica* MAGIC populations [46,57]. According to the recent genetic diversity analysis based on whole-genome sequencing data of 3,010 rice accessions [38], ten of the 16 Global MAGIC parents are *indica* rice accessions, which could explain the common genetic control of seed-related traits between our genetic material and other *indica* multi-parent populations. Also, deletions within the *GSE5* gene underlying the *GW5* locus are involved in the divergence between wide *japonica* grains and slender *indica* seeds [24], two ecotypes represented in the Global MAGIC population. This locus, along with the marker-trait associations found at the beginning of chromosome 5, fell within an extended region with large LD blocks, where about 50% of the sites are private alleles from the parent IAC 165 [31].

Interestingly, we detected a large genomic region on chromosome 12 controlling seed weight in the WW environment and ellipse eccentricity under drought stress. This region co-localizes with a large effect QTL, *qDTY_12.1_,* controlling grain yield under reproductive and seedling stage drought stress [58]. This QTL is stable across different genetic backgrounds and versatile to various environments and cropping systems [58,59]. It increases lateral root growth, transpiration efficiency, secondary panicle branching, and the number of filled grains per panicle [42]. Our results demonstrate that *qDTY_12.1_* could also play an important role in yield components under seedling stage drought.

Within the *qDTY_12.1_* region, we observe that the gene LOC_Os12g27810 carries a 15-nt deletion and a linked SNP that change the protein sequence, donated by Jinbubyeo, the only temperate *japonica* parent of our population. The MAGIC lines with the minor haplotype showed lower values of ellipse eccentricity (rounder seeds) than those with the major haplotype, and this increase in seed roundness is greater under drought (**Fig 8**), suggesting mechanisms of drought avoidance or tolerance [60]. Dixit et al. [42] found multiple intra-QTL genes underlying *qDTY_12.1_* and report that, while LOC_Os12g27810 is up-regulated under drought stress, it is not a putative functional partner of *OsNAM_12.1_.,* the main candidate gene underpinning the functionality of *qDTY_12.1_.* This study evaluated the response to drought stress in many traits but not grain size among near isogenic lines (NILs) with *qDTY_12.1_* donated from the *aus* variety Way Rarem into the *indica* variety Vendana. Given the relative frequencies of LOC_Os12g27810 haplotypes we observe in these subpopulations in the 3K genotypes, it is probable that our minor haplotype was not part of this previous study. While the LOC_Os12g27810 haplotypes segregating in the Global MAGIC population are good candidate alleles for functional differences underlying grain size and shape, it is possible that this locus acts independently from the previously described *qDTY_12.1_,* or that it is in only LD with the causal locus, scenarios that future studies can clarify. With our GBS site datasets we are investigating a small proportion of the genome-wide genetic variation within our MAGIC population; thus, the chances of detecting true trait causal variants are relatively small. If LOC_Os12g27810 is indeed causal, we were fortunate that it is also directly adjacent to a GBS cut site. Finally, we note that even if the minor haplotype could be a candidate for increasing yield components under drought, its ultimate performance will depend upon consumer acceptance of rounder seeds, which varies across different geographical areas of rice production and consumption [7].

Beyond determining the true involvement of LOC_Os12g27810 in grain shape under drought stress, our results clearly indicate a GxE component for ellipse eccentricity at this locus (**Fig 7**). We see non-random segregation of the SNP at S12_16392557 among the *indica* and *japonica* subpopulations identified by Wang et al. [38], and a suggestive evidence that the deletion is similarly segregating among subpopulations. The prevalence of the minor allele T (a nonsynonymous substitution) in the GJ-tmp and GJ-sbtrp groups might indicate different histories of environmental adaptation compared to the *indica* varieties, cultivated mostly in tropical regions. During its spread into northern, more temperate regions, *japonica* rice developed mechanisms of adaptation to colder climates [61] and long-day conditions [62]. Our results suggest possible adaptive responses to water deficit as well, and the availability of whole-genome sequences data for numerous and diverse Asian rice accessions offers opportunities for future, in-depth investigation of local adaptation to drought in rice.

We found evidence of GxE interactions for all the other trait-loci associations identified, with most allele effects being conditionally neutral (**Fig 8-S11**). This scenario, where an allele has a strong effect in one environment and a minimal or neutral effect in the contrasting environment, can contribute to local adaptation at the organism level [63]. Conditionally neutral alleles can also facilitate the development of rice cultivars that produce high yield under dry-seeding and early-stage drought stress, and with stable yield potential also in normally irrigated field conditions, through pyramiding of alleles with positive effects in each environment. On the contrary, it will be difficult to incorporate the few alleles with antagonistic effects on seed weight under drought and well-watered conditions; in this case, a beneficial allele in one environment might cause losses in yield under different conditions of soil moisture.

Our results underscore the importance of understanding the contribution of GxE interactions on trait variation better, which will be crucial in defining breeding goals and strategies for the development of varieties with either high and stable yield potential in different environmental conditions, or adapted to specific climates [64]. Recently, Diouf et al. [64] studied the genetic control of the responses of various traits to multiple abiotic stresses (*i.e.,* water deficit, salinity, and heat stress) in a tomato MAGIC population. They observed that genetic mean effects and phenotypic plasticity have both common and independent genetic control, a discovery that will allow to design efficient breeding strategies for the generation of tomato cultivars resilient to multiple stresses. In addition, they confirmed the crucial role of MAGIC populations in the study of trait variation under abiotic stress conditions. In rice, the Global MAGIC population that integrates both *indica* and *japonica* genetic background represent a powerful resource to investigate the effect of a combination of stresses to yield-related traits, and quantify the contribution of GxE interactions to trait phenotypic variation. Integration of high-throughput phenotyping with a diverse genetic material suited for multi-environmental trial is the answer to adapt rice cultivation to current and future climate fluctuations.

## Acknowledgments

We are thankful to Hei Leung and his group at IRRI for developing the rice Global MAGIC population. We thank Amelia Henry and her group also at IRRI for conducting field trials, sharing germplasm, and comments on an earlier version of this manuscript. We also acknowledge Noah Fahlgren and Haley Schuhl from the Donald Danforth Plant Science Center for their assistance in developing the PlantCV pipeline, and Daniela Urbina for helping with the seed phenotyping.

## Author contributions

Conceptualization: BTM.

Data curation: AM, BTM.

Formal analysis: AM.

Funding acquisition: BTM.

Investigation: AM.

Methodology: AM, BTM.

Project administration: AM, BTM.

Resources: BTM.

Software: AM, BTM.

Supervision: BTM.

Validation: AM.

Visualization: AM.

Writing – original draft: AM.

Writing – review & editing: AM, BTM.

## Supplementary captions

**Fig S1.** Principle steps of the PlantCV pipeline used to process all seed scans.

**Fig S2.** Principal component analysis of the Global MAGIC population and the 14 phenotyped parents, performed using the genetic profiles at the 7,795 GBS variants discovered on the Nipponbare reference genome. Parents’ name abbreviations: CYPR_S = Cypress; FEDE_0 = Fedearroz 50; IAC_5 = IAC 165; IET_0 = IET 14720; INIA_I = Inia Tacuari; IR_4_1 = IR 45427-2B-2-2B-1; IR_4_3 = IR 4630-22-2-5-1-3; IR_7_3 = IR 77186-122-2-2-3; IR_7_0 = IR 77298-14-1-2-10; IR_8__ = IR 84196-32 SAMBA MAHSURI(SUB1); JINB_O = Jynbubyeo; PSB_2 = PSB RC82; SANH_2 = SANHUANGZHAN NO. 2; WAB_5 = WAB 56-125

**Fig S3.** Correlation between trait measures of the 53 samples chosen randomly within the drought experiment. CH = convex hull.

**Fig S4.** Correlation between trait measures of the 54 samples chosen randomly within the well-watered experiment. CH = convex hull.

**Fig S5.** Graphical description of how the PlantCV algorithm estimates seed length, seed width, ellipse major and minor axis, and ellipse angle. The measures of seed length and width, along with ellipse angle, depends on the orientation of the rice seed on the scanner, causing inconsistencies across assays for the same genotype.

**Fig S6.** Distribution of the Global MAGIC parents and lines grown under drought stress.

**Fig S7.** Trait distribution in the Global MAGIC parents and lines grown under well-water conditions.

**Fig S8.** Seed area of the 14 phenotyped MAGIC parents under drought and well-watered conditions. Parent labels are colored according to the classification of Wang et. al 2018: black = *indica;* red = *japonica;* blue= basmati-type.

**Fig S9.** Ellipse major axis of the 14 phenotyped MAGIC parents under drought and well-watered conditions. Parent labels are colored according to the classification of Wang et. al 2018: black = *indica;* red = *japonica;* blue= basmati-type.

**Fig S10.** GWAS analysis for all analyzed traits using the Nipponbare site dataset. The dashed line indicates the threshold of significance FDR > 0.05.

**Fig S11.** GWAS analysis for all analyzed traits using the 93-11 site dataset. The dashed line indicates the threshold of significance FDR > 0.05.

**Fig S12.** Reaction norms of the standardized allele effects for all 93-11 sites significantly associated (FDR < 0.05) in at least one environment (or treatment) to ellipse eccentricity, solidity, and seed weight. Major allele trait effects standardized to trait standard deviation in each environment (y-axis) are plotted against the trait mean in each environment for the 243 lines common to the two environments (x-axis).

**Fig S13.** Reaction norms of the standardized allele effects for the Nipponbare site S5_1530423 significantly associated to ellipse eccentricity, convex hull vertices, and ellipse minor axis. The major allele trait effect standardized to trait standard deviation in each environment (y-axis) is plotted against the trait mean in each environment for the 243 lines common to the two environments (x-axis). The same reaction norms were observed for the 93-11 marker S5_1530423.

**Table S1.** Description of the 12 traits measured per seed using a user-designed PlantCV pipeline.

**Table S2.** Test of phenotypic differences between treatments (drought stress vs well-watered) and genotypes for the Gaussian traits using two-way analysis of variance (ANOVA).

**Table S3.** Test of phenotypic differences between treatments (drought stress vs well-watered) and genotypes for the non-Gaussian traits using Kruskal-Wallis test.

**Table S4.** Summary statistics of the ten measured traits between drought treated (DS) and well-watered (WW) samples.

**Table S5.** Eleven top ranked samples for seed area under drought stress.

**Table S6.** Significant genome-wide associations (FDR < 0.05) found in Nipponbare and 93-11. The associations are organized in intervals based on their proximity and overlap across genomes. The full lists of markers-trait associations for Nipponbare and 93-11 are reported in Table S7 and S8, respectively.

Table S7. Significant marker-trait associations (FDR < 0.05) found on the Nipponbare reference genome. Markers in bold have an antagonistic allelic effect under drought and well-watered conditions. FDR = False-Discovery-Rate; MAF = Minor Allele Frequency; SD = Standard Deviation.

**Table S8.** Significant marker-trait associations (FDR < 0.05) found on the 93-11 reference genome. FDR = False-Discovery-Rate; MAF = Minor Allele Frequency; SD = Standard Deviation.

**Table S9.** List of known QTL for drought tolerance and seed traits found in the surrounding regions (-/+ 2 MB) of the significant marker-trait associations. Source: SNP-Seek2.

**Table S10.** List of known genes for drought tolerance and seed traits found in the surrounding regions (-/+ 2 MB) of the significant marker-trait associations. Source: SNP-Seek2.

**Table S11.** List of known genes for drought tolerance and seed traits found in the surrounding regions (-/+ 2 MB) of the significant marker-trait associations. Source: Oryzabase.

**Table S12.** List of candidate genes found within the 4 MB windows drawn around the most significant associated markers.

**Table S13.** Haplotypes formed by the 16 sites of the gene LOC_Os12g27810 within the 3K accessions resequenced by Wang et al. [38].

**Table S14**. Frequency of the seven main haplotypes (present in more than one individual) among the nine rice subpopulation found by Wang et al. [38]. XI-1A = indica samples from East Asia; XI-1B = modern indica varieties of diverse origins; XI-2 = indica samples from South Asia; XI-3 = indica samples from Southeast Asia); GJ-tmp = East Asian temperate japonica samples; GJ-sbtrp = japonica accessions; GJ-trp = Southeast Asian tropical japonica samples; cA = Aus varieties; cB = Basmati and Sadri aromatic varieties.

